# Investigating Factors Driving Shifts in Subtype Dominance within H5Nx Clade 2.3.4.4b High-Pathogenicity Avian Influenza viruses

**DOI:** 10.1101/2025.04.23.650244

**Authors:** Elizabeth Billington, Cecilia Di Genova, Caroline J. Warren, Saumya S. Thomas, Simon Johnson, Sofia Riccio, Dilhani De Silva, Jacob Peers-Dent, Nigel Temperton, Kelly da Costa, Alexander M. P. Byrne, Maisie Airey, Audra-Lynne Schlachter, Jiayun Yang, Alejandro Nunez, Munir Iqbal, Marek J. Slomka, Ian H. Brown, Ashley C. Banyard, Joe James

## Abstract

H5Nx clade 2.3.4.4b high-pathogenicity avian influenza viruses (HPAIVs) have decimated wild bird and poultry populations globally since the autumn of 2020. In the United Kingdom (UK) and in continental Europe, the H5N8 subtype predominated during the first epizootic wave of 2020/21, with few detections of H5N1. However, during the second (2021/22) and third (2022/23) epizootic waves, H5N1 was the dominant subtype. The rapid shift in dominance from H5N8 to H5N1 was likely driven by a combination of virological, immunological, and/or host-related factors. In this study, we compared viral fitness and immunological responses in ducks, a key reservoir species, using dominant genotypes of H5N1 (genotype AB) and H5N8 (genotype A) from the second wave. While viral shedding dynamics were similar for both viruses, H5N8 was more pathogenic. Antigenic analysis of post-infection duck sera revealed that the haemagglutinin (HA) protein was antigenically similar across clade 2.3.4.4b H5 HPAIVs, but neuraminidase (NA) proteins displayed different patterns of cross-reactivity. We also modelled a scenario where ducks were pre-exposed to H5N1 (genotype C) or H5N8 (genotype A) from the first wave and subsequently challenged with either homologous or heterologous subtypes from the second wave (genotype AB or A). Despite the absence of seroconversion, pre-exposure to different subtypes resulted in varying clinical outcomes following challenge. These findings indicate that both viral and immunological factors likely played significant roles in the emergence and spread of H5Nx HPAIVs in wild bird populations.

## INTRODUCTION

Influenza A viruses are subtyped according to the genetic and antigenic relatedness of the two surface glycoproteins, haemagglutinin (HA) and neuraminidase (NA). Currently, seventeen different HA subtypes and 9 different NA subtypes of avian influenza virus (AIV) have been detected in wild birds. The HA protein can be broadly split into five clades, which form two groups; group one (clade 1, H8, H9, H12, H19; clade 2, H1, H2, H5 and H6; clade 3, H11, H13, and H16) and group two (clade 1, H3, H4, and H14; clade 2, H7, H10, and H15) [1, 2]. A similar approach can be applied to the NA protein, which can be divided into two groups (group one, N1, N4, N8 and N5; group two, N6, N9, N7, N2 and N3) [2]. Due to the segmented nature of the AIV genome, which consists of eight genomic segments, different HA and NA combinations can arise via reassortment when two or more AIVs co-infect the same cell.

Reassortment of the internal gene segments can also generate new genotypes with different biological properties. Wild aquatic birds (mainly Anseriformes [waterfowl; ducks and geese etc.], and Charadriiformes [shorebirds; gulls and waders etc.]) serve as the reservoir for AIV in which nearly every combination of HA and NA subtypes can be found [2].

Since 2014, there have been repeated introductions of clade 2.3.4.4 H5Nx high pathogenicity avian influenza virus (HPAIV) of the A/goose/Guangdong/1/96 (gsGd) lineage into the European poultry sector, driven by the movements of wild bird species on migratory pathways [3]. The gsGd lineage epidemiology has been recently dominated by clade 2.3.4.4b HPAIVs, which continue to circulate in wild bird populations, and because of continual reassortment with other circulating AIVs, multiple different NA subtypes have emerged and established dominance in wild birds across Europe with resultant poultry incursions, including H5N8 (2014/15 and 2016/17), H5N6 (2017), H5N8 (2020/21) and H5N1 (2021/22 and 2022/23) [4–10]. During winter 2020/21 (wave 1), the magnitude of H5Nx epizootics escalated considerably [11].

In the United Kingdom (UK) and continental Europe, the first H5Nx epizootic wave (2020/21 autumn/winter) was the largest ever at the time. Wave 1 was dominated by a single subtype, H5N8, which was responsible for 96% of detections in UK poultry. A minor population of the H5N1 subtype was also circulating during this period, with H5N1 being responsible for 4% of detections in UK poultry [9, 10].

During wave 1, all H5N8 HPAIVs detected in the UK shared >98.1% sequence identity across all gene segments existing as a single genotype (European Reference Laboratory [EURL]: genotype A; H5N8-W1 [12]), similarly all H5N1 detections also formed a single genotype (genotype C; H5N1-W1) [10].

At the end of the first wave (summer 2021) and at the start of the second wave (W2; 2021/22), H5N1 HPAIV emerged as the dominant subtype in the UK, with less than 1% of detections in wild birds and poultry being of other subtypes (H5N2 and H5N8), with only a single detection of H5N8 from a mute swan (*Cygnus olor*) in November 2021 [10, 13]. This initial H5N1 was genetically highly similar to H5N1-W1 which circulated as a minority subtype in the previous season (both were genotype C). Similarly, the single detection of H5N8 (H5N8-W2) was highly genetically related to the H5N8-W1 virus that had dominated previously (both were genotype A) [10].

As the second wave developed (autumn 2022), and transitioned into the third wave (2022/23), reassortment with other circulating AIVs contributed to increased genetic diversity within the H5N1 HPAIVs. However, importantly, little variation was observed in the HA gene between all H5N1 and H5N8 genotypes detected since wave 1. Four distinct H5N1 genotypes were detected more than once in poultry in the UK until the end of wave 3 (October 2023): genotypes AB, BB, C, and AL [10, 14–18]. These genotypes have had varied detection frequencies, with the AB genotype (H5N1-W2) providing 50.9% of detections since it was first detected at the start of the second wave in the UK. This genotype of H5N1-W2 shared the PB1, NP, NA, MP, and NS segments with H5N1-W1 [10, 19], but the PB2 and PA segments were most closely related to H5N3 AIVs from the first wave [10].

The rapid transition in dominance from H5N8 to H5N1 between wave 1 and wave 2 suggests (i) a potential phenotypic change in the virus, or (ii) potential differential immune evasion mechanisms between the subtypes. Therefore, in this study, virological and host factors which may influence the continued evolution of H5Nx clade 2.3.4.4b HPAIVs in Anseriformes were investigated. Specifically, we investigated the relationship between the two epizootic seasons, where H5N8 dominated in wave 1 and H5N1 dominated in wave 2. Using virus isolates representative of those circulating in wave 2 of the epizootic in 2021/22 (H5N1-W2 and H5N8-W2), we compared *in vivo* fitness in ducks and interrogated antigenicity against HA and NA. The changing subtype dominance was modelled by exposing ducks to wave 1 viruses (H5N1-W1 and H5N8-W1) and then challenging with homologous or heterologous wave 2 viruses (H5N1-W2 or H5N8-W2).

## METHODS

### Viruses and propagation

Six avian influenza virus (AIV) strains were used in this study (Table S1). Two viruses represented the first wave epizootic (Oct 2020–Sept 2021): A/chicken/England/030786/2020 (H5N8, genotype A; H5N8-W1; GISAID: EPI_ISL_17212363) and A/mute swan/England/234255/2020 (H5N1, genotype C; H5N1-W1; GISAID: EPI_ISL_766876). De-engineered versions of both viruses were generated by substituting the multi-basic cleavage site (MBCS) with a single-basic cleavage site (SBCS) via reverse genetics (RG) (described previously [20]), resulting in SB-H5N8-W1 and SB-H5N1-W1.

For in vivo challenge studies, two HPAIVs from the second wave epizootic (Oct 2021–Sept 2022) were used: A/chicken/Scotland/054477/2021 (H5N1, genotype AB; H5N1-W2; GISAID: EPI_ISL_9012696) and A/mute swan/England/298902/2021 (H5N8, genotype A; H5N8-W2; GISAID: EPI_ISL_13369742).

All viruses were propagated in 9-day-old specified pathogen free (SPF) embryonated fowl eggs (EFEs) and titrated in EFEs to determine the 50% egg infectious dose (EID_50_), as previously described [21]. The full genomes of all viruses used in this study were sequenced, as previously described [10] to confirm that no amino-acid polymorphisms had emerged following passage, when compared to the original sequence.

### Animals

High health status Pekin ducks (*Anas platyrhynchos*) (Cherry Valley hybrid; Cherry Valley Farms Ltd., UK) of mixed sex (52% female; 48% male, total 36) were obtained at one-day old and reared until 3 weeks of age at time of infection. Ducks were acclimatised for seven days prior to the first procedures which included swabbing (oropharyngeal (Op) and cloacal (C)) and bleeding from a superficial vein prior to inoculation. These samples were tested using an M-gene real time reverse transcription polymerase chain reaction (RT-PCR) [22] and serological testing [23] using the ID Screen® Influenza A Antibody Competition Multi-species ELISA kit (IDVet, France). For the IVPI test, white Leghorn SPF chickens (*Gallus gallus domesticus*) (Valo, Germany) were reared at APHA until 4 weeks-of-age.

### Experimental design

The statutory IVPI test was performed on de-engineered viruses (SB-H5N1-W1 and SB-H5N8-W1) as per WOAH guidelines [23]. Virus isolates were diluted 1:10 in isotonic saline, and 100 µl (10^8^ EID_50_/bird) was administered intravenously into the wing vein of 4-week-old chickens (n=10).

For infection and prior-immunity studies, two groups of immunologically naïve 3-week-old ducks (n=6/group) were inoculated via the ocular-nasal route with 10^5^ EID_50_ of either H5N1-W2 or H5N8-W2 in 100 µl of phosphate buffered saline (PBS) (Fig. S1). Oropharyngeal (Op) and cloacal (C) swabs were collected daily from 1–14 days post infection (dpi). Birds were euthanised upon reaching humane endpoints or at 14 dpi, when blood was collected via cardiac puncture under terminal anaesthesia.

For prior-exposure studies, two groups of ducks (n=12/group) were inoculated oronasally with 10^7^ EID_50_ of SB-H5N1-W1 (N1) or SB-H5N8-W1 (N8) (Fig. S1). At 13 dpi, blood was collected from all birds, after which each group was split into two (n=6/group) and challenged oronasally with 10^5^ EID[[ of either H5N1-W2 or H5N8-W2. This formed four groups: N1/N1 and N8/N1 (H5N1-W2 challenge), and N1/N8 and N8/N8 (H5N8-W2 challenge). From 1–14 days post challenge (dpc), Op and C swabs were collected daily. Birds were euthanised at humane endpoints or at 14 dpc (28 dpi total), when terminal blood collection was undertaken.

### Clinical scoring and monitoring

Birds were observed and scored against a defined clinical score sheet (Table S2) twice a day (morning and afternoon). Following the development of clinical disease, birds were additionally observed at a third timepoint (evening). Birds were terminated whenever clinical endpoints were reached (cumulative score >7; Table S2). For ethical reasons, any birds housed singularly, due to euthanasia or death of others within the group, were also euthanised.

### Clinical sample collection, processing, and immunohistochemistry

Swabs were individually cut and placed into 1ml of Leibovitz L-15 Medium (Gibco; (LM) [24]), and the supernatant used for RNA extraction or stored at −80°C until further use. During post-mortem examination, approximately 50 milligrams (mg) of tissue from selected organs was collected into 1ml LM and mechanically homogenised, with the clarified homogenate being used for RNA extraction. In addition, tissues were harvested from all major organs and fixed in 10% (v/v) neutral buffered formalin for a minimum period of 5 days before being embedded in paraffin. Four-micron thick serial sections were stained with haematoxylin and eosin (H&E), and for immunohistochemistry (IHC), using a mouse monoclonal anti-influenza A nucleoprotein antibody (Statens Serum Institute, Copenhagen, Denmark) as described previously [25]. The overall distribution of virus-specific staining in each tissue was assessed using a semi-quantitative scoring system (0 = no staining, 1 = minimal, 2 = mild, 3 = moderate and 4 = widespread staining) modified from [26]. Specificity of immunolabelling was assessed in positive control sections by replacing the primary antibody with a matching mouse IgG isotype; no non-specific cross-linking was observed.

### Environmental sample collection and processing

Following infection with HPAIV at 14 dpi, environmental samples of drinking water and pond water were collected on alternate days from each duck pen. Air samples were collected, as described previously [24, 27], from the environment of ducks infected with either H5N1-W2 or H5N8-W2. Viral RNA (vRNA) was extracted directly from water samples, and from supernatants of gelatine air filters dissolved in LM as described previously [27].

### RNA extraction and AIV reverse transcription Real-Time PCR (RT-PCR)

vRNA was extracted from LM obtained from swabs, tissues, feathers, and environmental samples using the MagMAX^TM^ CORE Nucleic Acid Purification Kit (Thermo Fisher Scientific™) and the robotic Mechanical Lysis Module (KingFisher Flex system; Life Technologies), according to the manufacturer’s instructions. Extracted RNA were tested by the M-gene RT-PCR using the primers and probes as previously described [22]. A ten-fold dilution series of RNA extracted from a stock of H5N1-W2 HPAIV with a known EID_50_ titre was used to construct a standard curve and Ct values converted to relative equivalent units (REU) as previously described [24, 28]. RT-PCR Ct values <36 (>10^1.794^ REU) were considered as AIV positive, Ct values ≥36 (≤10^1.794^ REU) were considered negative.

### Serum processing and hemagglutination inhibition assays (HI)

Whole blood samples were centrifuged for 5 minutes at 2000 rpm to separate the serum from the clot. Sera were incubated at 56°C for 30 minutes to inactivate complement. The duck sera were pre-absorbed by adding packed chicken red blood cells and incubated at 4°C overnight to prevent non-specific agglutination [23]. Hemagglutination inhibition assays (HI) were conducted using different antigens initiated at four HA units. Reciprocal HI titres of >1/16 were considered seropositive as defined internationally [23]. All sera were also tested by the ID Screen® Influenza A Antibody Competition Multi-species ELISA kit (IDVET, France) according to the manufacturer’s instructions.

### Influenza pseudotype virus (PV) generation

Lentiviral pseudotype viruses (PVs) containing H5Nx glycoproteins were generated for H5N1-W2 (genotype C) (A/chicken/England/053052/2021), H5N1-W2 (genotype AB) (A/chicken/Scotland/054477/2021) and H5N8-W1 (genotype A) (A/chicken/England/030786/2020) (Table S1). Existing PV were used for H5N8-14 (A/gyrfalcon/Washington/41088-6/2014) [29]. The HA and NA genes were subcloned from pHW2000 into pCAGGS using a directional *NotI-XhoI* restriction digest strategy. PVs were generated as previously described [29] employing p8.91 gag-pol vector as retroviral core. PV titres were assessed by transduction of HEK293T/17 cells as previously described [29] employing the Bright-Glo assay system (Promega) and read as relative luminescence per unit (RLU).

### Pseudotype virus neutralisation assay (PVNA)

For PVNA, serum was first prediluted 1:100 and tested in duplicate against H5 PVs. Chicken reference serum raised against for A/chicken/Scotland/59 (H5N1) (product code: RAA7002) and A/duck/England/036254/14 (H5N8) (Product code: RBB6425) (APHA Scientific, UK). Briefly, sera were serially diluted 2-fold in a 96-well white plate and 10^6^ RLU of PV was added to each well. The plate was incubated for 1 hour at 37[ to allow binding of the antibody to the antigen. Next, 10^4^ HEK293T/17 cells were added to each well. PV-only and cell-only controls were included in the plate to represent the 0% and 100% neutralisation of the PV. The plate was incubated for 48h at 37[ at 5% CO_2_ before reading.

Data were normalised and plotted on a neutralisation percentage scale and the reciprocal of the serum dilution which induces 50% neutralisation or IC_50_ was calculated as previously described [30].

### Pseudotype-based enzyme-linked lectin assay (pELLA)

To investigate NA reactive antibodies HA and NA PVs were employed. H5Nx PVs were generated and titrated via the pELLA method as previously described [31]. Optical density at 450nm (OD_450_) was determined using the Multiskan™ FC microplate photometer with SkanIt™ Software for data analysis. Readings were normalised to 100% and 0% OD_450_, and the dilution that resulted in 90% OD_450_ was selected for inhibition assays. The inhibition of NA activity by serum samples was evaluated via pELLA. Serum was tested in duplicate against H5N1 and H5N8 PVs. Sera were initially diluted 1:10 and then serially diluted 2-fold in 50µL SD. 50µL of these dilutions were then transferred to fetuin-coated plates containing 50µL of SD to all wells, excluding SD-only and PV-only controls, representing 100% and 0% inhibition, respectively. 50µL of the H5Nx PV 90% OD_450_ as determined in the titration was then transferred to all wells of the fetuin-coated plate except for SD-only control. All other steps were followed as per NA PV titration via the pELLA. The IC_50_ was calculated as the inverse dilution of serum that resulted in 50% inhibition of NA activity as determined via GraphPad Prism.

### Statistical analysis

Area under the curve (AUC) analysis of individual bird shedding profiles was performed using GraphPad Prism v8, with values expressed as REU. Birds that did not shed were assigned an AUC value of 0 REU. Mean AUC values were calculated and compared for statistical significance using one-way ANOVA with multiple comparisons in GraphPad Prism v8. Statistical significance was defined as p < 0.05. For pELLA analysis, mean and standard deviation medians were derived for each data point based on two replicates per dilution. Kaplan–Meier survival curves were evaluated using the Gehan–Breslow–Wilcoxon test in GraphPad Prism v8.

## RESULTS

### Comparative fitness in ducks of 2.3.4.4b H5N1 and H5N8 HPAIV circulating during the second epizootic wave

The *in vivo* fitness of H5N1-W2, a HPAIV representative of the dominant genotype (AB) during the second epizootic wave, was compared with H5N8-W2 (genotype A) from the same time period (Fig. S1). Inoculation of naïve ducks with either virus resulted in productive infection, with high vRNA shedding from the Op cavity (Fig. 1A and B). Both viruses showed similar shedding profiles; however, H5N8-W2 exhibited a higher mean peak shedding on 3 dpi, while H5N1-W2 peaked later, at 4 dpi, with a lower mean titre. Notably, H5N1-W2-infected ducks shed vRNA for a longer duration. C shedding was broadly similar, though a higher proportion of ducks shed vRNA after H5N1-W2 infection (H5N1-W2, 100%; H5N8-W2, 50%). However, AUC analysis showed no significant difference between H5N1-W2 and H5N8-W2 in overall Op or C shedding (Op, p-value 0.229; C: p-value 0.477). Environmental contamination was found in the drinking and pond water with vRNA detected from 3 dpi in both groups, but less consistently in the H5N1-W2 group (Fig. 1C and D). vRNA was present in water on 3, 9, and 11 dpi for H5N1-W2, and on 3, 5, 7, and 9 dpi for H5N8-W2. Airborne vRNA was only detected at 7 dpi in the H5N1-W2 group.

**Fig. 1.**
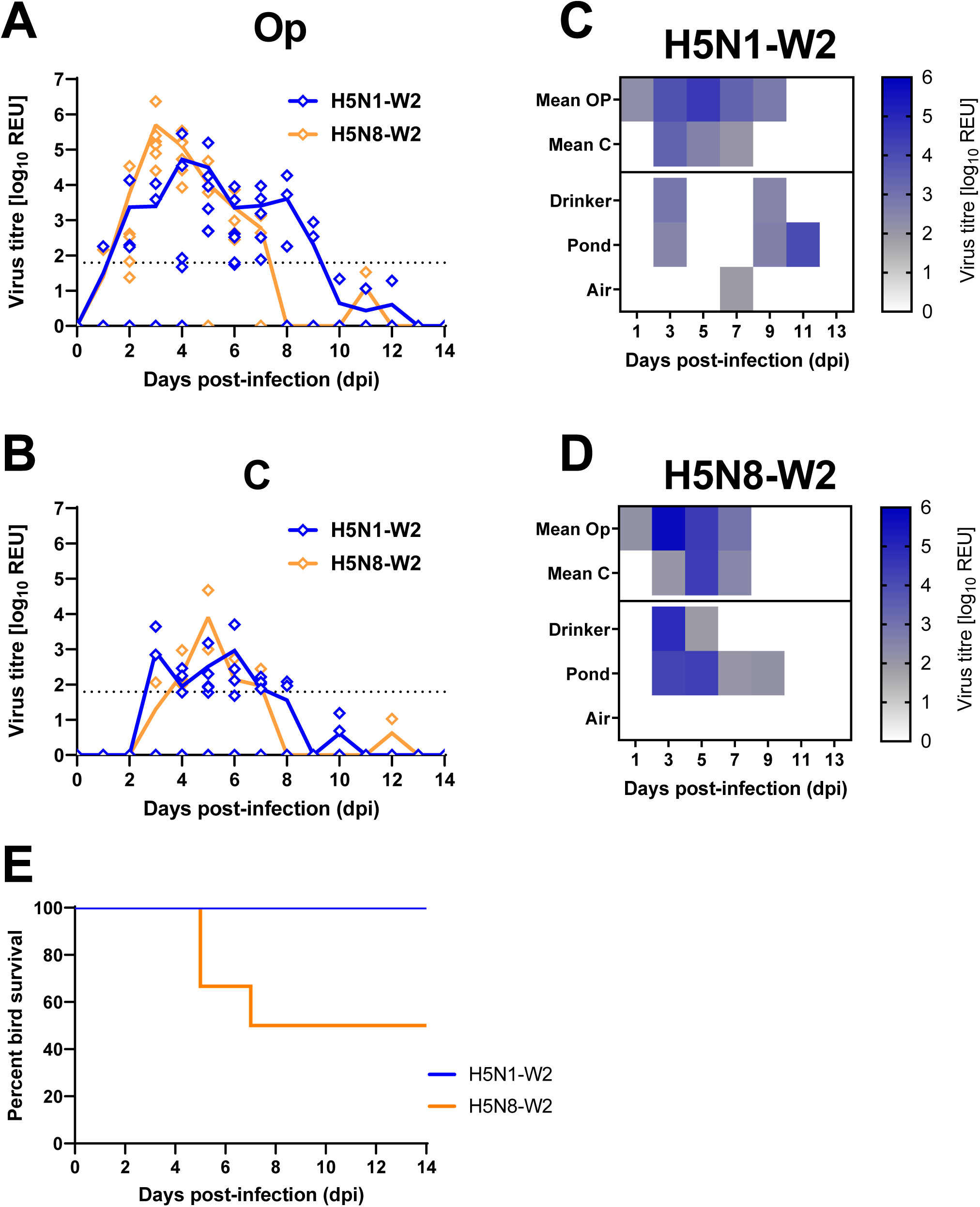
Virus shedding, environmental contamination, and survival from ducks infected with H5N1-W2 and H5N8-W2 HPAIVs. **A and B**. Viral shedding measured from swab samples collected from the oropharyngeal (Op; A) and cloacal (C; B) cavities of naïve ducks following infection with H5N1-W2 (blue) or H5N8-W2 (orange) HPAIV. Viral RNA (vRNA) titres were quantified by M-gene RT-PCR and expressed as log_10_ relative expression units (REU). Individual data points are shown, with coloured lines indicating the group mean. The dotted horizontal line represents the diagnostic Ct threshold (Ct 36.00; 10^1.794^ REU) for M-gene RT-PCR positivity. **C and D**. Heatmaps representing quantification of vRNA (log_10_ REU) for H5N1-W2 (**C**) and H5N8-W2 (**D**). The top two rows of each panel correspond to the mean shedding from OP and cloacal C swabs, as shown in panels A and B. Environmental samples (drinker water, pond water, and air) were collected on alternate days. Heatmap shading indicates vRNA detection: white denotes no Ct value, grey represents samples below diagnostic threshold (10^1.794^ REU), and blue indicates positive samples. **E.** Percentage survival of ducks over a 14-day period post-infection (dpi) with H5N1-W2 (blue) or H5N8-W2 (orange) HPAIV.

### Clinical outcomes in ducks with H5N1 and H5N8 HPAIV circulating during the second epizootic wave

All ducks infected with H5N1-W2 survived until the end of the study (14 dpi), whereas only 50% of those infected with H5N8-W2 survived (two ducks were euthanised at 5 dpi and one at 7 dpi) (Fig. 1E). H5N1-W2 infection induced fewer clinical signs, with a total clinical score of 8, compared to a score of 90 in the H5N8-W2 group (Table S2). Ducks infected with H5N8-W2 also exhibited more severe neurological signs, including tremors (7 counts), torticollis (1 count), and seizures (1 count) (Table S2, Fig. S2A and B).

Post-mortem examination was conducted on three H5N1-W2-infected ducks and six H5N8-W2-infected ducks. As no H5N1-W2 ducks required culling, comparisons between groups are only possible at 14 dpi. Among the three H5N8-W2-infected ducks culled between 5–7 dpi (ducks 101, 103, and 104), vRNA was detected in all sampled tissues from ducks 103 and 104 (5 dpi), while duck 101 tested negative in the heart, lung, caecal tonsil, proventriculus, and liver (Fig. 2A). Notably, all three showed high vRNA levels in brain and feather samples, with mean titres of 10^8.41^ and 10^5.48^ REU, respectively. Of the six surviving ducks culled at 14 dpi, vRNA was detected only sporadically, and at a low level, in brain, muscle, or feathers in one H5N8-W2-infected duck (106) and one H5N1-W2-infected duck (100). Another H5N1-W2-infected duck (99) had detectable vRNA only in the pancreas and kidney.

**Fig. 2.**
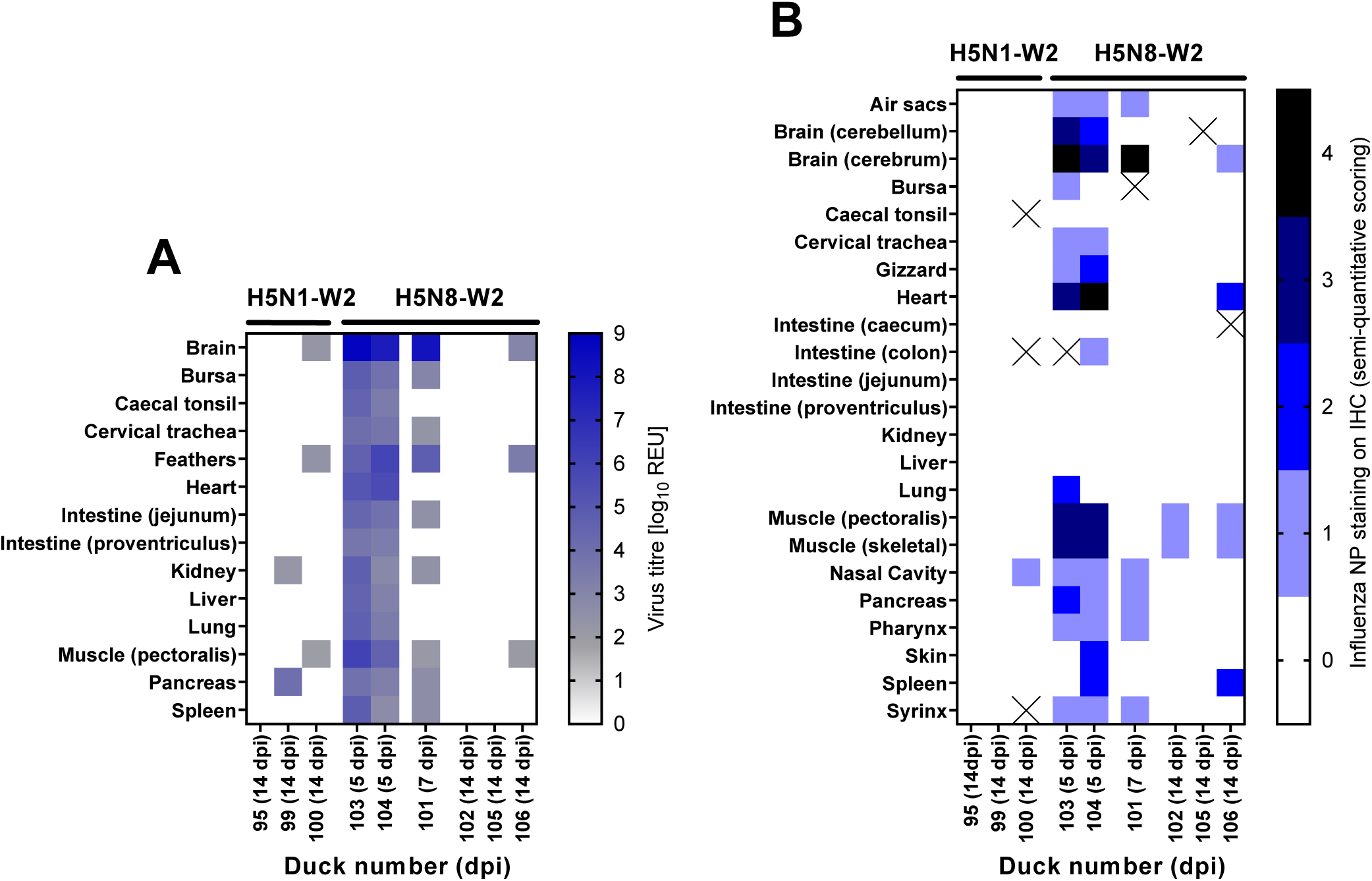
Virus distribution in tissues of ducks infected with H5N1-W2 or H5N8-W2 HPAIV. Quantification of viral RNA (vRNA) in tissue samples collected from nine ducks infected with H5N1-W2 (n = 3) or H5N8-W2 (n = 6) HPAIV. Ducks were euthanised due to severe clinical signs or culled at the study endpoint (14 days post-infection, dpi). vRNA levels were measured by M-gene RT-PCR and expressed as log_10_ relative expression units (REU), represented as a heatmap. Samples with Ct values below the diagnostic threshold (10^1.794^ REU) are shown in grey, positive samples in blue, and samples with no detectable Ct value in white. **B.** Semi-quantitative scoring of anti-NP staining intensities and distribution in immunohistochemistry (IHC) analysis of tissues (lower to higher intensities, 0-4), while the “X” indicates that no sample was available for IHC.

IHC analyses was also performed on tissues from these birds. At 5 dpi, brain samples from H5N8-W2 ducks showed acute non-suppurative encephalitis with extensive viral antigen in neurons (Fig. 2B and S3A and B). By 14 dpi, only minimal multifocal gliosis and sparse antigen were observed, indicating viral clearance. In the heart, acute myocardial degeneration with monocytic infiltration and abundant antigen was seen at 4–5 dpi. Pectoral muscle displayed mild non-suppurative myositis with variable antigen labelling. By 14 dpi, lesions in both heart and muscle had resolved to mild chronic inflammation and lymphoid hyperplasia. Minimal antigen remained in macrophages within lymphoid aggregates, suggesting resolution of active viral replication.

### Serological responses and antigenic cross-reactivity between H5N1 and H5N8 subtypes

Serological responses following infection with H5N1-W2 and H5N8-W2 were compared. All surviving infected ducks (n=6 H5N1-W2; n=3 H5N8-W2) were positive by NP ELISA (data not shown) and seroconverted against the homologous antigens in the HI assay, at 14 dpi (Fig. 3A). Lower, but not statistically significant, HI titres were detected when sera from both infections were tested against the H5N8-W2 compared to the H5N1-W2 antigen. No statistically significant differences were observed between reactivity of viruses with sera of the same subtype from wave 2 or wave 1 (H5N1-W2 compared to H5N1-W1 and the reciprocal) (Fig. 3A). To assess HA-specific antibody responses, PVs displaying only the HA proteins of clade 2.3.4.4b H5N1 and H5N8 strains were used, alongside an earlier clade 2.3.4.4c H5 (H5N8-14) PV as a reference. Although both sera neutralised H5N8-14 PV, IC_50_ values were significantly lower than those for the more recent H5 PVs (Fig. 3B). No significant differences in neutralisation titres were found between the 2.3.4.4b H5 PVs using either serum (Fig. 3B). To evaluate NA-specific responses, pELLA was performed using PVs containing both HA and NA. H5N1 and H5N8 antisera inhibited their homologous PVs with IC_50_ values of 10^3.68^ and 10^4.89^, respectively (Fig. 3C and D). Pre-infection sera showed no cross-reactivity to either PV. Post-infection sera from both groups (H5N1-W2 and H5N8-W2 sera) robustly inhibited H5N8 PV (IC_50_ 10^4.28^ for H5N1-W2 and 10^4.58^ for H5N8-W2). However, H5N1-W2 sera only inhibited the H5N1 PV (IC_50_ 10^3.68^). H5N8-W2 sera did not inhibit H5N1 PV, with inhibition curves failing to reach 50% (Fig. 3C and D).

**Fig. 3.**
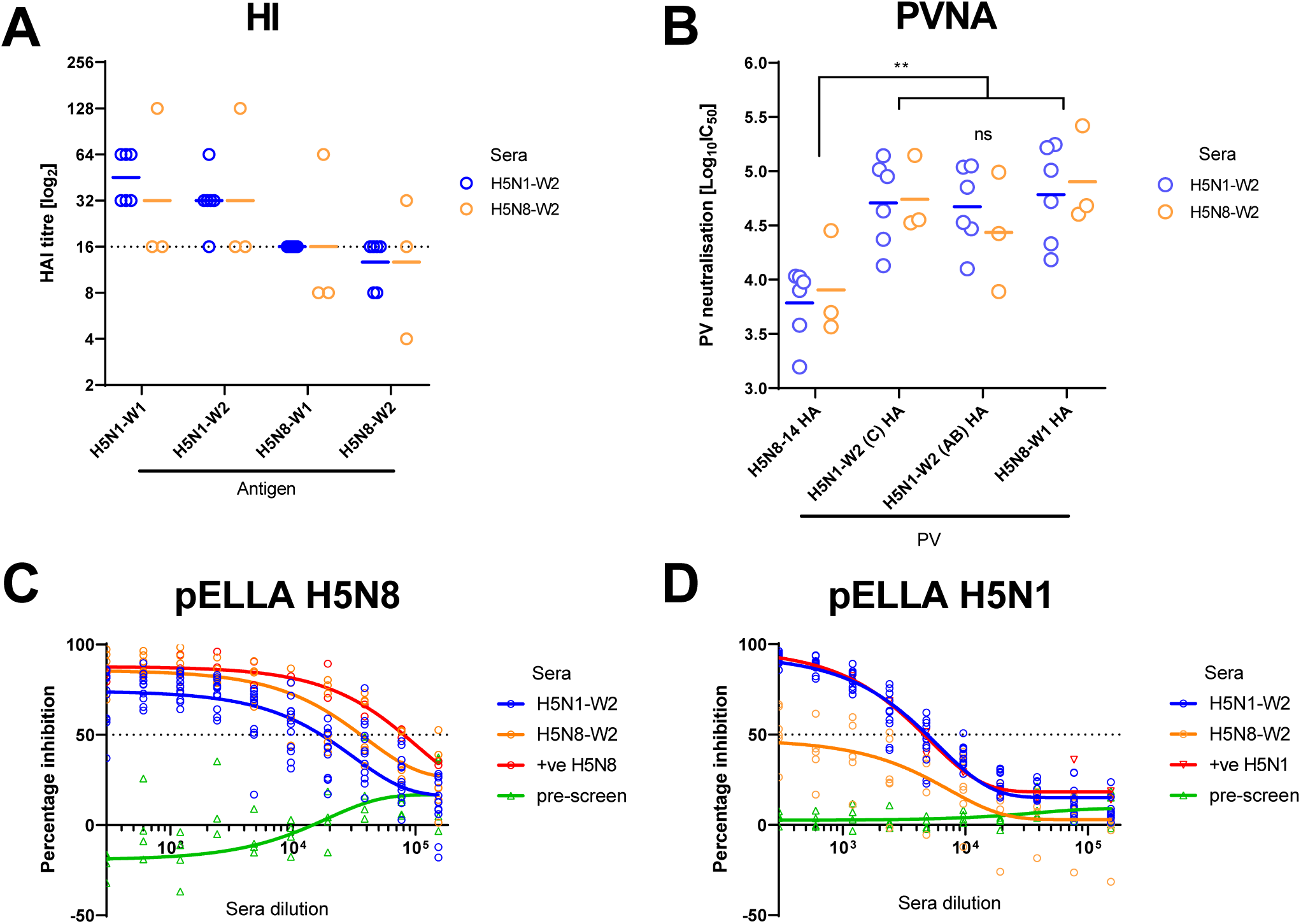
Serological responses and antigenic characterization of H5N1-W2 and H5N8-W2. Serological responses using sera collected from ducks at 14 dpi infected with H5N1-W2 AB (blue) or H5N8-W2 A (orange); sera assessed using HI (**A**) or lentiviral pseudotype viruses (PVs) containing H5Nx influenza virus HA or HA and NA glycoproteins (**B-D**). Coloured horizontal bars indicate geometric mean titres, while the dotted line indicates the diagnostic threshold for positivity. **A**. HI using sera from ducks infected with H5N1-W2 or H5N8-W2, against antigens with homologous or heterologous NAs from 2021 or 2020. **B.** PV neutralisation titres (inhibition concentration 50%; IC_50_) using PVs containing H5 HA only evaluated against individual sera from H5N1-W2 or H5N8-W2 infected ducks. (**C-D**) Sera assessed by pELLA using PVs containing HA and NA glycoproteins tested against pre-infection (green), homologous subtype positive control (red), H5N1-W2 (blue) or H5N8-W2 (orange) tested against post-infection sera from H5N1-W2 or H5N8-W2 infected ducks.

### H5N1 and H5N8 clade 2.3.4.4b H5Nx HPAIV without MBCSs have reduced pathogenicity yet are able to productively infect chickens

To replicate the natural exposure scenario observed between 2020–2022, we next modelled sequential infection in wild birds, with initial exposure to wave 1 viruses (H5N1-W1 or H5N8-W1) followed by re-infection with wave 2 viruses (H5N1-W2 or H5N8-W2). To ensure survival after primary exposure, we attempted to reduce the pathogenicity of the viruses while retaining the antigenicity. Therefore, we used RG to generate low pathogenic variants (SB-H5N1-W1 and SB-H5N8-W1) by replacing the multi-basic cleavage site (MBCS) with a single-basic cleavage site (SBCS) (Fig. S4C). Both SB viruses reached high titres in embryonated fowl eggs (SB-H5N1-W1, 10^10.50^ EID_50_/ml; SB-H5N8-W1, 10^10.00^ EID_50_/ml), comparable to their wildtype HPAIV counterparts (H5N1-W1, 10^9.85^; H5N8-W1, 10^9.48^). IVPI testing confirmed a low pathogenic phenotype (SB-H5N1-W1, 0.15; SB-H5N8-W1, 0.0), yet all surviving birds seroconverted by 10 dpi, confirmed by homologous HI titres (Fig. S4A). One mortality occurred in the IVPI at 6 dpi in the SB-H5N1-W1 group. vRNA was detected in multiple tissues and swabs from this animal (Fig. S4B) but sequencing confirmed retention of the engineered SBCS (Fig. S4C).

### The effect of homo- and hetero-subtypic prior exposure on shedding dynamics of the H5N1 and H5N8 clade 2.3.4.4b HPAIV

We next inoculated ducks, as surrogates for wild aquatic birds, with SB-H5N1-W1 or SB-H5N8-W1 viruses, followed by homologous (N1/N1, N8/N8) or heterologous (N1/N8, N8/N1) HPAIV challenge with H5N1-W2 or H5N8-W2 (Fig. S1). Following inoculation with the de-engineered viruses, vRNA was detected in only three of 12 ducks in the SB-H5N1-W1 groups (N1/N1, N1/N8), all below the positive threshold; no vRNA was detected in the SB-H5N8-W1 groups (N8/N8, N8/N1). No clinical signs were observed, and all ducks were seronegative by HI at 13 dpi (HI titres <2).

Following challenge with H5N1-W2, ducks previously exposed to SB-H5N1-W1 (N1/N1) exhibited a 4-day delay in vRNA detection in Op swabs compared to naïve controls (Fig. 4A). A similar delay was observed in C swabs (Fig. 4E). Despite this, both groups reached comparable peak vRNA levels, with Op approximately 2 log_10_ higher than C shedding (Fig. 4A, E). Total shedding (AUC) did not differ significantly between groups (Op, p-value 0.9991; C, p-value 0.9941). vRNA was detected in water from the N1/N1 group from 7–11 dpi; a 4-day delay compared to the naïve H5N1-W2 group (Fig. 5A vs Fig. 1C). In the heterologous challenge (N1/N8), prior SB-H5N1-W1 exposure delayed shedding onset by 2 days relative to naïve H5N8-W2-infected ducks, though both groups peaked at 3 dpi (Fig. 4B). Water contamination was detected at 3 dpi in both groups but persisted longer (until 9 dpi) in the naïve (H5N8-W2) group (Figs. 1D, 5D). The delay in shedding onset in the N1/N1 and N1/N8 groups was most apparent early in infection (Fig. 4A–B).

**Fig. 4.**
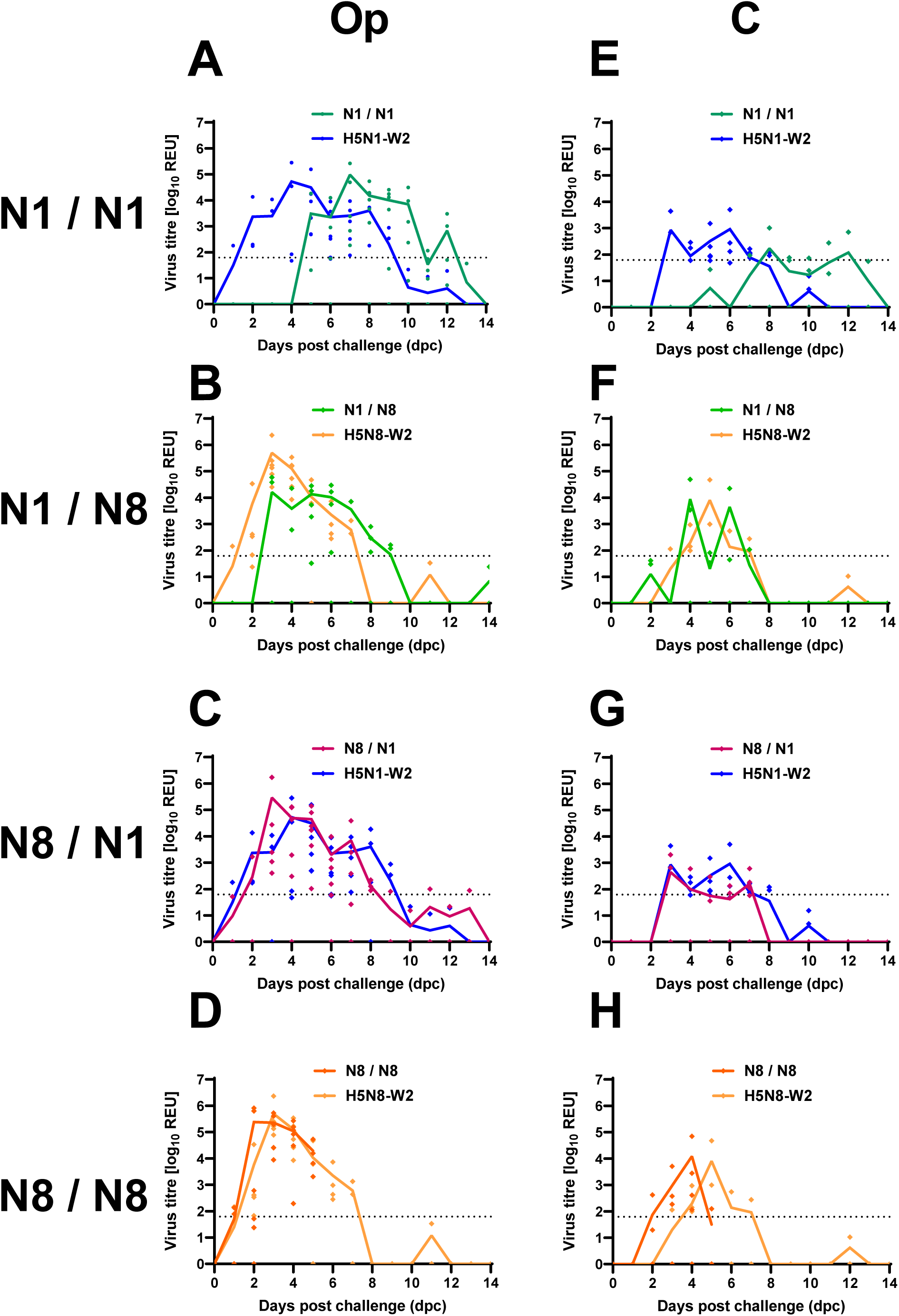
Virus shedding from ducks infected with H5N1-W2 and H5N8-W2 HPAIV with prior exposure to de-engineered H5N1-W1 and H5N8-W1 AIVs. Groups of ducks were initially inoculated with either H5N1-W1 (N1) or H5N8-W1 (N8) de-engineered AIVs. After 14 days post-inoculation (dpi) the groups were divided, and ducks were challenged with either HPAIV H5N1-W2 or H5N8-W2. This resulted in homologous (N1/N1 **(A and E)** and N8/N8 **(D and H)**, or heterologous (N1/N8 **(B and F)** and N8/N1 **(C and G)** subtype combinations. Shedding was measured from oropharyngeal (OP) (**A-D**) and cloacal (C) (**E-H**) swabs following challenge. Shedding from naïve birds also shown for reference. vRNA titres were determined by M-gene RT-PCR and expressed as log_10_ REU. Individual shedding values are shown by coloured symbols, with group mean shedding shown by continuous coloured lines. Dotted horizontal lines indicate the M-gene RT-PCR positive cut-off (10^1.794^ REU).

**Fig. 5.**
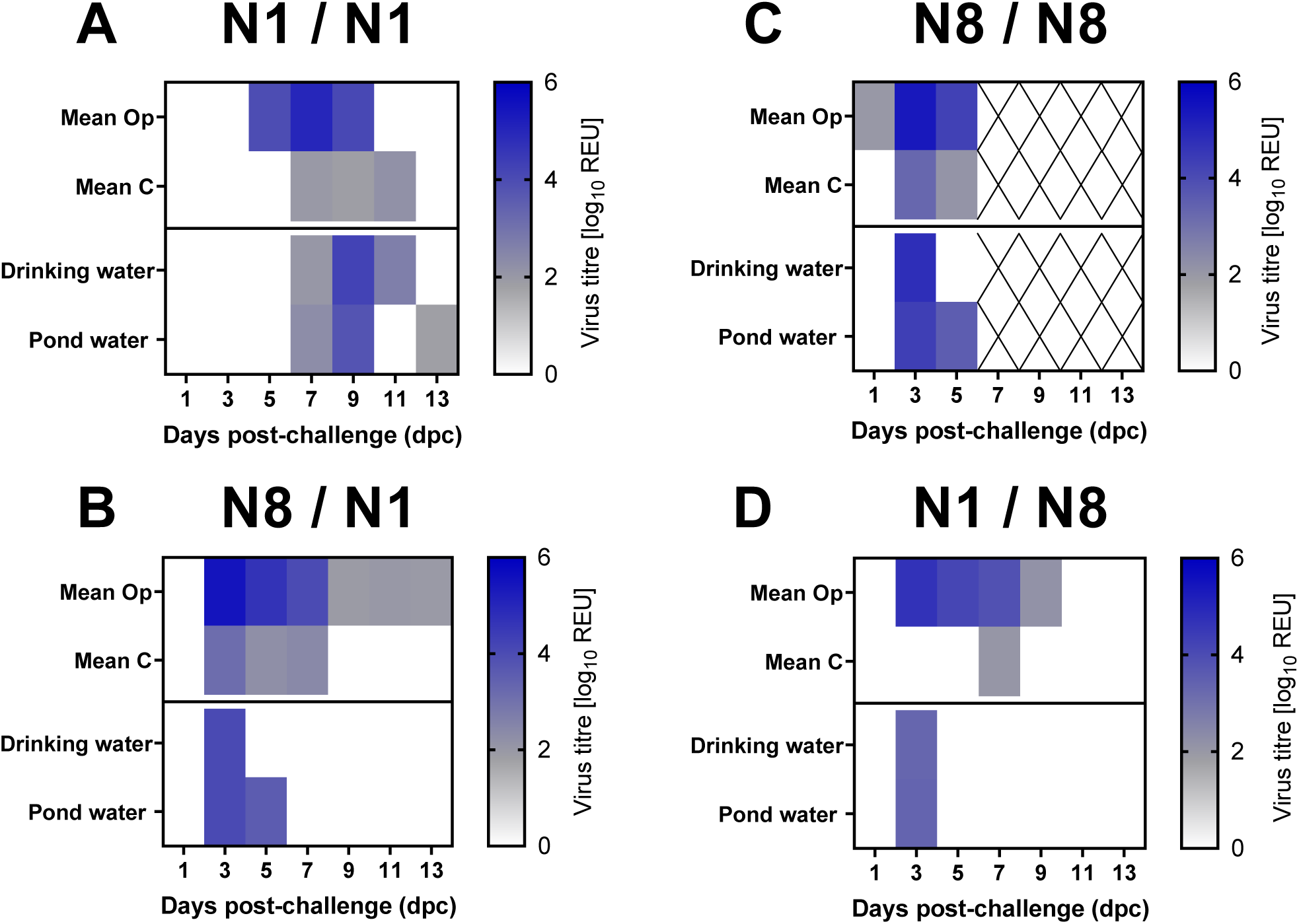
Viral environmental contamination from ducks challenged with H5N1-W2 and H5N8-W2 HPAIV, following prior exposure to de-engineered H5N1-W1 and H5N8-W1 AIVs. Environmental specimens were collected between 1-13 dpc from drinker and pond waters of ducks. The ducks had been challenged with HPAIV H5N1-W2 or H5N8-W2 but previously been exposed to homologous (N1/N1 **(A)** and N8/N8 **(C)**) or heterologous (N1/N8 **(D)** and N8/N1 **(B)**) subtype of de-engineered H5N1-W1 or H5N8-W1 AIVs. vRNA titres were determined by the M-gene RT-PCR. Samples shown as a heatmap with no Ct in white, below the diagnostic cut-off (10^1.794^ REU) shown in grey, positive samples in blue. ‘X’ indicates that no sample was collected at the specific time point. The top two rows on all graphs represent the mean oropharyngeal (OP) and cloacal (C) shedding titres from all birds.

In contrast, prior SB-H5N8-W1 exposure had minimal impact on vRNA shedding. Ducks subsequently challenged with H5N1-W2 (N8/N1) or H5N8-W2 (N8/N8) showed similar vRNA shedding profiles to naïve controls (Fig. 4C–D, G–H). Environmental contamination was also similar (Figs. 5B–C). Water from the N8/N1 group showed comparable levels to naïve H5N1-W2-infected ducks, although shedding was more prolonged in the naïve H5N1-W2 ducks (Figs. 1C, 5B). For the N8/N8 group, vRNA was detected in water, but sample collection ceased at 5 dpi due to mortality in all ducks, limiting comparison (Figs. 1D, 5C, 6B).

**Fig. 6.**
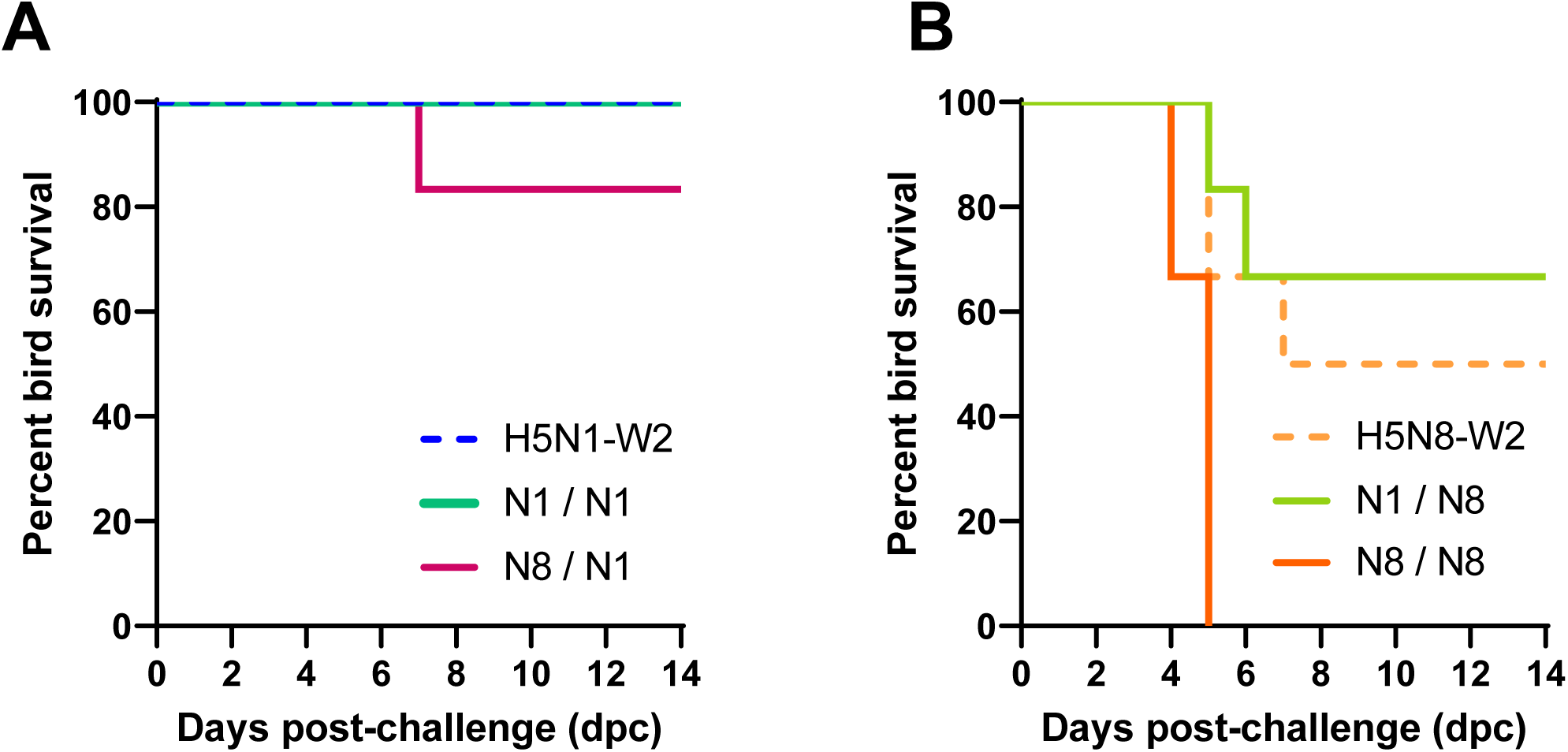
Survival of ducks challenged with H5N1-W2 and H5N8-W2 HPAIV, with or without prior exposure to de-engineered H5N1-W1 and H5N8-W1 AIVs. Percentage survival of ducks until 14 dpc following challenge with (A) HPAIV H5N1-W2 or (B) H5N8-W2 with no prior exposure (hashed lines), or ducks which had previously been exposed to (solid lines) homologous (N1/N1 and N8/N8) or heterologous (N1/N8 and N8/N1) subtypes of de-engineered H5N1-W1 or H5N8-W1 AIVs.

### The effect of homo- and hetero-subtypic prior exposure on disease outcomes and pathogenesis

In the H5N1-W1 pre-exposed ducks, no mortality was observed following challenge with H5N1-W2 (N1/N1), consistent with the survival of naïve H5N1-W2-infected ducks (Fig. 6A). Both groups also exhibited reduced clinical severity compared to ducks exposed to or infected with H5N8 viruses (Fig. S2). In contrast, prior exposure to H5N1-W1 reduced mortality and clinical signs following H5N8-W2 challenge (N1/N8) compared to the naïve group, but not statistically significantly (Fig. 6B, Table S2).

However, in the N8 pre-exposed group, homologous H5N8 re-challenge (N8/N8) resulted in statistically significantly higher mortality (p = 0.0139, Gehan-Breslow-Wilcoxon test) and greater clinical severity compared to the naïve H5N8 group (Fig. 6B and S2B, D), with 100% of ducks culled due to severe disease (two at 4 dpc, four at 5 dpc), compared to 50% in the naïve group (two at 5 dpc, one at 7 dpc) (Fig. 6). Prior H5N8-W1 exposure also increased mortality upon H5N1-W2 challenge (N8/N1).

Tissue tropism was assessed only in ducks that succumbed to infection. vRNA was detected in multiple tissues from N1/N8, N8/N1, and N8/N8 groups, with the highest titres in brain samples (Fig. 7A). High vRNA levels were also found in feathers, cervical trachea, and intestines of N1/N8 and N8/N8 ducks. In contrast, no vRNA was detected in tissues from the N8/N1 bird, which also showed lower IHC scores across tissues compared to H5N8-W2-challenged birds (Fig. 7A and B). Histopathological lesions and immunolabelling in N1/N8 and N8/N8 ducks mirrored those in naïve H5N8-W2-infected birds (Figs. S3C, S3D). Moderate multifocal non-suppurative encephalitis with gliosis and perivascular cuffing was observed in the brain, with abundant neuronal antigen. Acute myocardial degeneration and necrosis with co-localised viral antigen were evident in the heart and pectoral muscle (Fig. 7B). In contrast, N8/N1 ducks displayed minimal lesions, and no tissue samples were available from the N1/N1 group.

**Fig. 7.**
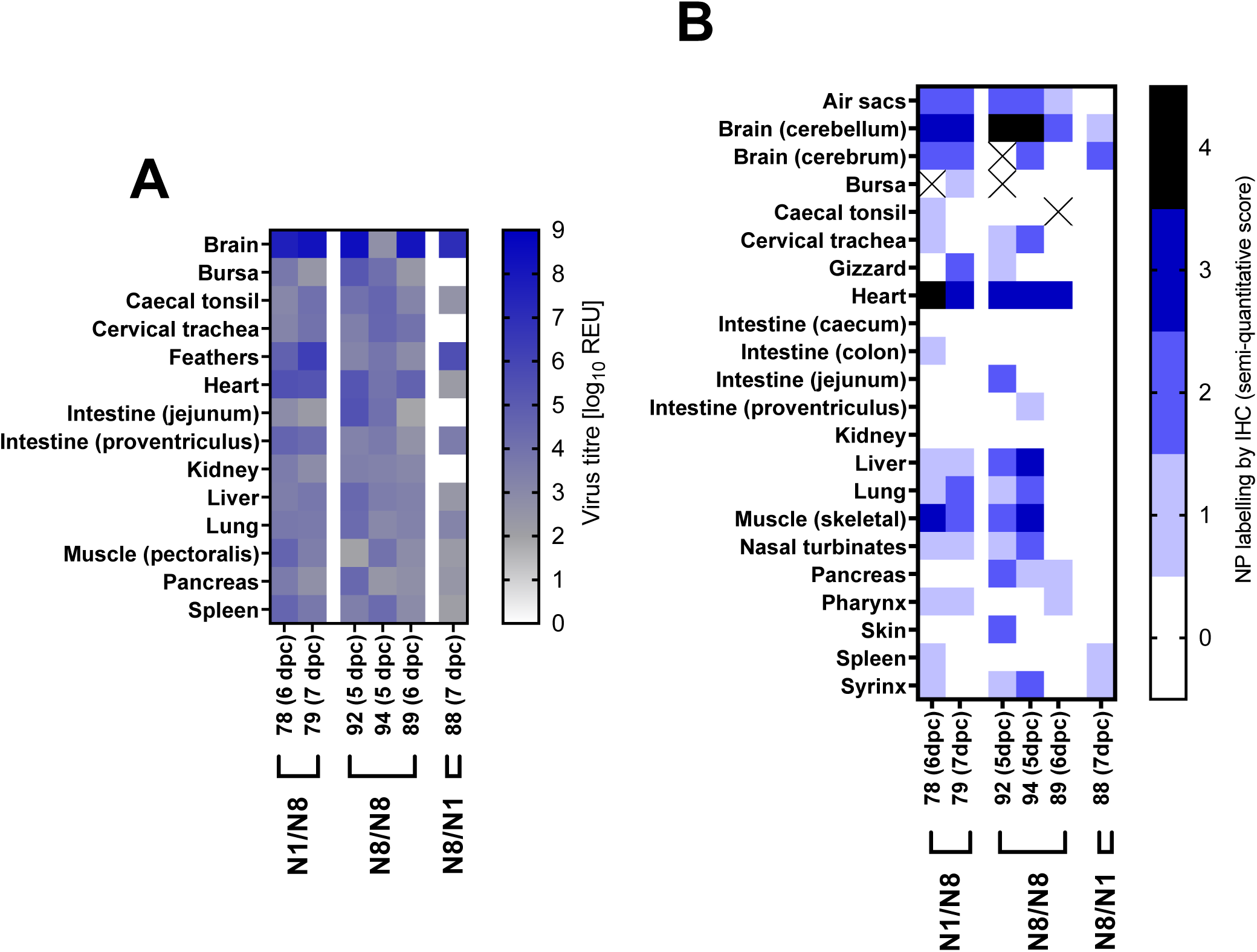
Organ tropism in ducks with prior exposure to de-engineered AIVs before H5N1-W2 and H5N8-W2 HPAIV challenge. Tissue samples were collected from six ducks following H5N1-W2 or H5N8-W2 HPAIV challenge, which had previously been exposed to a homologous (N1/N1 and N8/N8) or heterologous (N1/N8 and N8/N1) subtype of de-engineered H5N1-W1 or H5N8-W1 AIVs. All ducks were euthanised at indicated times (dpc) as the result of severe clinical signs. **A.** vRNA levels, measured by M-gene RT-PCR, expressed as log_10_ REU and represented as a heatmap. Samples below the diagnostic cut-off (10^1.794^ REU) shown in grey, positive samples in blue, and no Ct in white. Individual bird identifiers allow for comparisons to serological response graphs and IHC-staining results. **B.** Semi-quantitative scoring of anti-NP staining intensities and distribution in IHC analysis of tissues (lower to higher intensities, 0-4), demonstrating abundant viral antigen in the brain, heart and muscle following N1/N8 and N8/N8 challenge.

## DISCUSSION

During the H5Nx clade 2.3.4.4b panzootic, a notable shift in NA subtype dominance occurred in the UK and continental Europe. In the 2020–2021 season (wave 1), H5N8 predominated, with 26 infected poultry premises (IPs) and 285 wild bird cases reported in the UK, compared to only 2 IPs and 18 wild bird detections of H5N1. However, by 2021–2022 (wave 2), H5N1 became dominant, with 157 IPs and 1,629 wild bird detections, while H5N8 was detected only once, in a single mute swan (*C. olor*), with no poultry cases [9, 10]. This rapid shift may reflect (**i**) changes in viral virulence or infectivity, (**ii**) differential immune evasion, or (**iii**) a combination of both.

To assess virological fitness, representative H5N1 and H5N8 viruses from wave 2 were compared *in vivo*. Both H5N1-W2 and H5N8-W2 established productive infections, with all ducks shedding virus and seroconverting. Shedding kinetics were similar, with higher viral loads from Op swabs than C swabs. H5N1-W2 showed a more prolonged Op shedding period, though total viral load did not differ significantly between groups. Notably, H5N8-W2 caused 50% mortality, while H5N1-W2 caused none. Previous studies have shown that an earlier H5N1-W2 HPAIV (genotype C) had comparable infectivity and transmissibility to other clade 2.3.4.4 HPAIVs, with enhanced fitness in Anseriformes [32–34].

Recently, Bordes, Germeraad [35] compared two H5N1 genotypes of different dominance (genotype C [wave 1 – low] and genotype AB [wave 2 – high]) in three species of Anseriformes (Pekin ducks [*A. platyrhynchos*], Eurasian wigeon [*Mareca Penelope*] and barnacle geese [*Branta leucopsis*]) [35]. In all three species the dominant genotype AB was less pathogenic than less dominant genotype C; in pekin ducks no mortalities were seen for genotype AB compared to 40% mortalities with genotype C. The observation of high mortalities with early wave 1 H5N1 (genotype C) parallels with our findings of high mortalities for early wave 1 H5N8. Furthermore, consistent with our observations here, viral shedding kinetics were comparable between genotype AB and genotype C [35]. Bordes, Germeraad [35] did not assess transmission dynamics, and likewise transmission was not assessed here in our study. However, several studies highlighted a correlation between transmission efficiency and the extent of environmental contamination, particularly in water sources, for H5Nx 2.3.4.4b viruses [24, 33]. In our study, environmental contamination, particularly of pond and drinking water, was detected more frequently with H5N1-W2. This, combined with reduced vRNA shedding duration and higher mortality for H5N8-W2, suggests that H5N1-W2 may transmit more efficiently in wild waterfowl. The infrequent contact between wild birds implies that premature host death could limit transmission opportunities, as previously proposed [36]. Together, these data indicate phenotypic differences between H5N8 and H5N1 that may explain the ecological shift in subtype dominance from H5N8 in wave 1 to H5N1 in wave 2 in the UK and Europe.

Following initial characterisation of infection dynamics and pathogenicity, we investigated the potential role of antibody-mediated immunity in the observed shift in subtype prevalence between the first two outbreak waves. Some avian species are susceptible to HPAIV yet can survive infection. Ringing data suggest wild Anseriformes have life expectancies ranging from 3 to 20 years, depending on species [37], implying that individuals tolerant to infection may persist across multiple epizootics and encounter diverse AIV subtypes and genotypes [24, 38]. Antigenic relatedness and associated cross-reactivity between similar HA and/or NA proteins can complicate interpretation of serological data in wild birds.

Nevertheless, HA- and NA-specific antibodies have been detected in several wild waterfowl species [33, 39, 40]. To explore this further, we first assessed serological responses in naïve ducks surviving infection with H5N1-W2 or H5N8-W2, finding that all birds seroconverted by HI. Although not statistically significant, titres were lower when sera from both groups were evaluated against H5N8-W2 antigen than H5N1-W2. To better resolve HA-specific responses, we evaluated antibody titres using HA-only PVNAs. No significant differences in IC_50_ values were observed between sera from H5N1 and H5N8-infected birds. This similarity in HA antigenicity among clade 2.3.4.4b H5Nx viruses aligns with previous findings [41, 42] and likely reflects the high amino acid identity across their HA proteins (Table S3).

Although most of the antibody response in birds exposed to AIV targets HA [43], antibodies against NA also contribute to protective immunity [44, 45]. However, the composition and function of the antibody repertoire in wild bird species remain poorly characterised, particularly regarding the relative roles of HA- and NA-specific responses in protection. To investigate this, we assessed NA-specific antibodies using the pELLA assay [31]. While pELLA does not directly quantify neutralisation, it correlates well with NA inhibition activity [46]. Sera from ducks surviving infection showed detectable NA-specific antibodies, indicating a reactive immune response to NA [44, 45]. Previous studies have demonstrated a protective role for NA antibodies in Canada geese (*Branta canadensis*) and mallards (*A. platyrhynchos*), where prior LPAIV exposure reduced disease severity upon NA-homologous H5Nx HPAIV challenge [47, 48]. Similarly, NA antibodies have provided protection against H5N1 HPAIV in chickens [49]. These findings support the concept of “permissive immunity”, whereby NA antibodies limit viral release from infected cells, thereby mitigating disease without fully blocking infection [50, 51], in line with our observations. Additionally, neutralising NA antibodies against N1 and N8 have been detected in wild ducks in North America [52]. Interestingly, H5N1-W2-specific antisera in our study exhibited broader cross-subtype reactivity, inhibiting both H5N1 and H5N8 PVs, while H5N8-W2 antisera reacted only with homologous PVs. This raises the hypothesis that NA-reactive antibodies may persist following H5Nx infection, with H5N1-induced antibodies conferring broader cross-reactivity.

Notably, this pattern mirrors the epidemiological shift from H5N8 to H5N1 dominance between waves 1 and 2, raising the possibility that immunity to H5N8 may have facilitated the emergence of H5N1 by allowing subsequent infection in previously exposed hosts.

To further examine the impact of prior subtype exposure on infection outcome, we de-engineered the MBCS of H5N1-W1 and H5N8-W1 HPAIVs to contain a SBCS, minimising potential severe outcomes ducks. The low pathogenicity phenotype of both mutants was confirmed by IVPI, as per international guidelines [21]. Unexpectedly, for ducks inoculated with 10^7^ EID_50_ of these de-engineered viruses no viral shedding or seroconversion was detected. This contrasted with chickens in the IVPI, which developed robust serological responses, including one mortality with vRNA detected in multiple tissues, while retaining the SBCS in HA. Additionally, *in ovo* amplification produced similar viral titres for both wildtype and de-engineered viruses, together suggesting the modified viruses remained replication competent in animal systems. Collectively, these findings indicate that while the viruses are viable, the MBCS is essential for productive infection of ducks with clade 2.3.4.4b H5Nx viruses.

Whilst the lack of seroconversion to the de-engineered viruses was unexpected, other host responses outside of antibody generation are also likely important for infection outcome following AIV infection, and so we investigated the effect of homosubtypic and heterosubtypic re-infection. Prior exposure had limited impact on vRNA shedding following HPAIV challenge. However, in the N1/N1 group, shedding and environmental contamination were delayed compared to naïve H5N1-W2-infected ducks, with fewer positive cloacal swabs. This delay was more pronounced in homologous (N1/N1) than heterologous (N1/N8) challenge (4 vs. 2 days, respectively). Disease severity also varied depending on the priming subtype. Prior SB-H5N1-W1 exposure reduced clinical scores and mortality following challenge with either H5N1-W2 or H5N8-W2. Conversely, prior SB-H5N8-W1 exposure increased disease severity and mortality for both challenge viruses. This may reflect differences in (**i**) infection efficiency of the challenge virus, (**ii**) replication of the SB-H5N8-W1 virus, or (**iii**) host immune modulation and the clonal nature of the priming vs. challenge viruses. Earlier studies found that pre-exposure to LPAIV, especially with seroconversion, conferred protection from HPAIV-induced disease [11, 53, 54], though these studies used older birds (13 weeks–18 months) compared to the 3-week-old ducks used here. Age-related immune differences, particularly within the first six weeks post-hatch [55–57], therefore may explain the differences observed. In the N8/N8 group, we obtained inverse results following challenge, with the pre-exposed birds having higher mortality than the naïve/N8 group. Unlike previous prior exposure studies that used genetically distinct LPAIVs [11, 54, 58], in the current study we used RG to de-engineer the MBCS and retain close genetic relationships between de-engineered AIV W1 exposure and HPAIV W2 challenge. A recent study exposing mallards to a H4N6 found minimal antibody responses, as we saw here, but detected evidence of complement activation early in infection [59]. We did not investigate complement activation, but this could be a factor to be considered in future studies of this type. While the exact mechanism remains uncertain, prior exposure does not greatly impact viral shedding but does appear to alter disease severity. While cross-subtype immunisation and challenge has been reported to cause vaccine-associated enhanced respiratory disease (VAERD) in pigs and ferrets [60, 61], there have been no reports of VAERD in avian hosts, but subtype mismatch during our prior exposure and challenge has shown an increase in clinical scores. When considering the epidemiological switch in dominance from H5N8 to H5N1, birds would potentially be exposed to H5N8 and subsequently encounter H5N1. In this scenario, our data suggest that prior exposure to H5N8 may not protect against subsequent exposure to H5N8, but could allow ducks to tolerate H5N1 infection, although allowing infection and transmission of H5N1 among Anseriformes. All N8/N1 ducks shed H5N1-W2 to similar titres as the H5N1-W2 naïve birds, but 80% survived infection.

In conclusion, this study identifies distinct virological and serological differences between H5N1-W2 and H5N8-W2 that may explain the rapid shift in subtype dominance during the 2021–2022 epizootic in the UK and continental Europe. While both viruses demonstrated comparable shedding kinetics and environmental contamination in infected ducks as surrogates for wild Anseriformes, H5N8-W2 exhibited higher pathogenicity, leading to greater mortality. However, the delayed onset of shedding and the increased survival associated with H5N1-W2 suggests that H5N1 may be more adept at sustaining transmission, particularly in wild bird populations. Furthermore, NA-reactive antibodies were detected following infection with both subtypes. Notably, N1 antisera cross-reacted with N8, but the reverse was not observed, suggesting that prior H5N8 exposure may not impede subsequent H5N1 infection at the wild waterfowl population scale. In addition, prior exposure may enhance the severity of disease outcome outside of humoral responses, though mechanisms are unknown. Together these data suggest that in ducks genotype A H5N8 from the second epizootic wave is more pathogenic than genotype AB H5N1 from the same period, and that H5N1 pathogenicity has decreased between epizootic waves, whereas H5N8 pathogenicity has been maintained or increased. These phenotypic characteristics could favour selection of H5N1 in a partially immune H5N8 population, but the ability of H5N1 viruses to at least in part evade host antibody responses to H5N8 could create an environment in the natural reservoir favouring their selection and persistence. These factors may have a direct correlation with the emergence and complete shift to H5N1 HPAIVs where previously H5N8 was dominant.

Following initial selection and shift, H5N1 rapidly underwent reassortment to produce a highly variable number of genotypes that also has likely contributed to the rapid dissemination and spread of this subtype [10]. H5N1 has continued to emerge, reassort and genetically evolve with LPAIVs present in wild birds, co-existing as multiple genotypes that through increasing opportunity for infection and fitness in multiple avian species, remain in dynamic circulation on a global scale. These insights underscore the complex interplay of viral fitness, host immunity, and environmental factors in shaping the dynamics of gsGD-lineage H5Nx HPAIV epizootics. Certainly, a more robust assessment of different HPAIV genotypes is required to understand the impact of these viruses on different avian populations.

## Author statements

### Ethics and Safety Statement

The *in vivo* study was reviewed and approved by the local Animal and Plant Health Agency (APHA) Animal Welfare and Ethical Review Body (AWERB) to comply with the relevant UK legislation, in accord with the UK Home Office (HO) Project License PP7633638. Material utilised for all experimentation was initially isolated in EFEs under HO license P5275AD31. According to the UK’s Advisory Committee on Dangerous Pathogens (ACDP) and the Specified Animal Pathogens Order (SAPO); the gsGd lineage H5Nx HPAIVs are classified as ACDP hazard group level three pathogens and SAPO level four, hence all animal and laboratory work involving infectious material was conducted within licensed containment level 3 facilities at APHA. Welfare monitoring of up to three times daily was undertaken to assess the birds for humane endpoints following the onset of clinical disease, enabling any decisions to be made concerning the need for euthanasia. All birds had access to food and water *ad libitum*.

## Acknowledgements

The authors would like to thank Nadia Chew, Amy Miller and Elena Mather for collection and processing of samples from the *in vivo* experiments, and the postmortem and histopathology teams in the Pathology department for collection and processing of tissues.

## Author contribution

EB, CW, ST, SJ, SR, AS, AN, JJ and MJS conducted the *in vivo* experimental work. CDG, EB and ST conducted serological work. JY and MI produced the viruses by reverse genetics. NT and KdC provided logistical support and training for PVNA and pELLA. EB, CW, AMPB, JJ, MJS and CDG analysed the data. EB, CW, CDG, MJS, IHB, ACB and JJ wrote the manuscript. JJ, ACB and IHB secured funding.

