## Supplementary Figures and Tables for "Investigating Factors Driving Shifts in Subtype Dominance within H5Nx Clade 2.3.4.4b High-Pathogenicity Avian Influenza viruses"

### Table S1

**Table S1.** Viruses used in this study.

| Abbreviation | Virus strain name | Description | EURL genotype | Subtype | GISAID accession number |
|---|---|---|---|---|---|
| H5N1-W1 | A/mute swan/England/234255/2020 | Wave 1 H5N1 | C | H5N1 | EPI_ISL_766876 |
| H5N1-W2 | A/chicken/Scotland/054477/2021 | Wave 2 H5N1 | AB | H5N1 | EPI_ISL_9012696 |
| H5N8-W1 | A/chicken/England/030786/2020 | Wave 1 H5N8 | A | H5N8 | EPI_ISL_17212363 |
| H5N8-W2 | A/mute swan/England/298902/2021 | Wave 2 H5N8 | A | H5N8 | EPI_ISL_13369742 |
| SB-H5N1-W1 | A/mute swan/England/234255/2020 | Wave 1 H5N1 with MBCS replaced with SBCS | C | H5N1 | EPI_ISL_766876* |
| SB-H5N8-W1 | A/chicken/England/030786/2020 | Wave 1 H5N8 with MBCS replaced with SBCS | A | H5N8 | EPI_ISL_17212363* |

**wild type* sequence; multi-basic cleavage site (MBCS); single basic cleave site (SBCS).

### Table S2

**Table S2.** Total counts of each clinical sign observed following infection with H5N1-21 or H5N8-21 with or without prior exposure.

| Clinical Sign | Individual clinical sign severity score (1-7) | Group | | | | | |
|---|---|---|---|---|---|---|---|
| | | **H5N1-W2** | **H5N8-W2** | **N1/N1** | **N1/N8** | **N8/N8** | **N8/N1** |
| Found dead | 12 | 0 | 0 | 0 | 0 | 0 | 0 |
| Euthanised | 11 | 0 | 3 | 0 | 2 | 6 | 1 |
| Changes in huddling | 1 | 0 | 7 | 3 | 2 | 5 | 4 |
| Eyes closed | 1 | 0 | 1 | 1 | 0 | 0 | 1 |
| Conjunctivitis | 1-2* | 4 | 1 | 10 | 4 | 20 | 36 |
| Changes in body position | 1 | 0 | 12 | 13 | 18 | 13 | 11 |
| Oedema | 2-3 | 0 | 0 | 0 | 0 | 0 | 0 |
| Cyanosis of extremities | 2-3* | 0 | 0 | 0 | 0 | 0 | 0 |
| Lethargy | 1 | 2 | 21 | 10 | 9 | 15 | 5 |
| Lack of engagement with enrichment | 1 | 0 | 22 | 9 | 5 | 5 | 2 |
| Oronasal discharge | 2 | 0 | 1 | 0 | 0 | 1 | 0 |
| Diarrhoea | 2-3* | 0 | 13 | 4 | 7 | 8 | 16 |
| Perceived weight reduction | 1-3* | 1 | 1 | 0 | 0 | 1 | 0 |
| Loss of balance | 2 | 0 | 2 | 0 | 0 | 6 | 2 |
| Tremors | 2-3* | 1 | 7 | 0 | 8 | 6 | 5 |
| Torticollis | 4 | 0 | 1 | 0 | 0 | 0 | 0 |
| Seizure | 7 | 0 | 1 | 0 | 2 | 1 | 0 |
| Paralysis | 7 | 0 | 0 | 0 | 0 | 0 | 1 |
| **Total** | | **8** | **90** | **50** | **55** | **81** | **83** |

* Range in score depends on severity of the clinical sign

### Table S3

| | | H5N8-W2 | H5N1-W2 | H5N1-W1 | H5N8-W1 |
|---|---|---|---|---|---|
| PB2 | H5N8-W2 | 100.00 | 98.16 | 98.68 | 99.47 |
| | H5N1-W2 | 98.16 | 100.00 | 98.94 | 97.89 |
| | H5N1-W1 | 98.68 | 98.94 | 100.00 | 98.41 |
| | H5N8-W1 | 99.47 | 97.89 | 98.41 | 100.00 |
| PB1 | H5N8-W2 | 100.00 | 98.55 | 98.21 | 99.74 |
| | H5N1-W2 | 98.55 | 100.00 | 99.04 | 98.28 |
| | H5N1-W1 | 98.21 | 99.04 | 100.00 | 98.21 |
| | H5N8-W1 | 99.74 | 98.28 | 98.21 | 100.00 |
| PA | H5N8-W2 | 100.00 | 98.46 | 98.10 | 99.72 |
| | H5N1-W2 | 98.46 | 100.00 | 99.12 | 98.74 |
| | H5N1-W1 | 98.10 | 99.12 | 100.00 | 98.39 |
| | H5N8-W1 | 99.72 | 98.74 | 98.39 | 100.00 |
| HA | H5N8-W2 | 100.00 | 99.29 | 99.46 | 99.65 |
| | H5N1-W2 | 99.29 | 100.00 | 99.46 | 99.65 |
| | H5N1-W1 | 99.46 | 99.46 | 100.00 | 99.82 |
| | H5N8-W1 | 99.65 | 99.65 | 99.82 | 100.00 |
| NP | H5N8-W2 | 100.00 | 97.99 | 97.93 | 99.00 |
| | H5N1-W2 | 97.99 | 100.00 | 98.96 | 98.39 |
| | H5N1-W1 | 97.93 | 98.96 | 100.00 | 98.96 |
| | H5N8-W1 | 99.00 | 98.39 | 98.96 | 100.00 |
| NA | H5N8-W2 | 100.00 | 30.94 | 30.29 | 97.95 |
| | H5N1-W2 | 30.94 | 100.00 | 96.64 | 30.02 |
| | H5N1-W1 | 30.29 | 96.64 | 100.00 | 29.32 |
| | H5N8-W1 | 97.95 | 30.02 | 29.32 | 100.00 |
| MP | H5N8-W2 | 100.00 | 98.11 | 99.02 | 99.37 |
| | H5N1-W2 | 98.11 | 100.00 | 98.37 | 98.74 |
| | H5N1-W1 | 99.02 | 98.37 | 100.00 | 99.67 |
| | H5N8-W1 | 99.37 | 98.74 | 99.67 | 100.00 |
| NS | H5N8-W2 | 100.00 | 86.10 | 88.00 | 98.84 |
| | H5N1-W2 | 86.10 | 100.00 | 97.63 | 86.43 |
| | H5N1-W1 | 88.00 | 97.63 | 100.00 | 88.35 |
| | H5N8-W1 | 98.84 | 86.43 | 88.35 | 100.00 |

H5N1-W1 (A/mute swan/England/234255/2020), EPI_ISL_766876; H5N8-W1 (A/chicken/England/030786/2020), EPI_ISL_17212363; H5N1-W2 (A/chicken/Scotland/054477/2021), EPI_ISL_9012696; H5N8-W2 (A/mute swan/England/298902/2021), EPI_ISL_13369742.

**Table S3**. Amino acid (aa) differences between different H5Nx HPAIVs using concatenated protein sequences

### Fig. S1

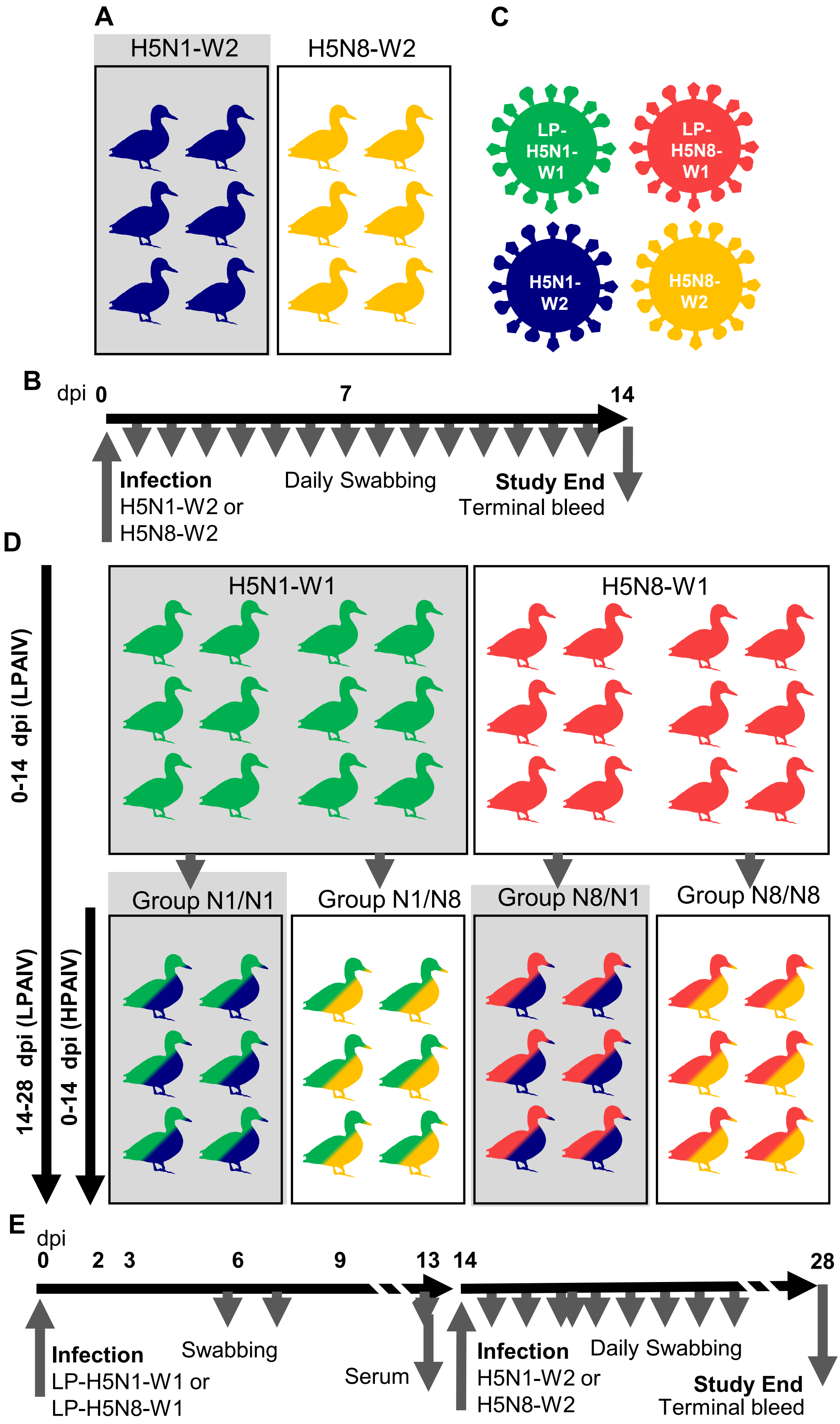

**Fig. S1. H5N1-W2 and H5N8-W2 *in vivo* experimental designs.**

**A and B**. Schematic diagram of the experimental design used to assess variation in fitness between H5N1-W2 and H5N8-W2 in naive ducks. **C**. Viruses used in this study. **D and E**. Schematic diagram of the experimental design used to assess infection outcomes from HPAIV H5N1-W2 or H5N8-W2 in ducks which had previously been inoculated with a homologous (N1/N1 and N8/N8) or heterologous (N1/N8 and N8/N1) subtype of de-engineered H5N1-W1 or H5N8-W1 AIVs.

### Fig. S2

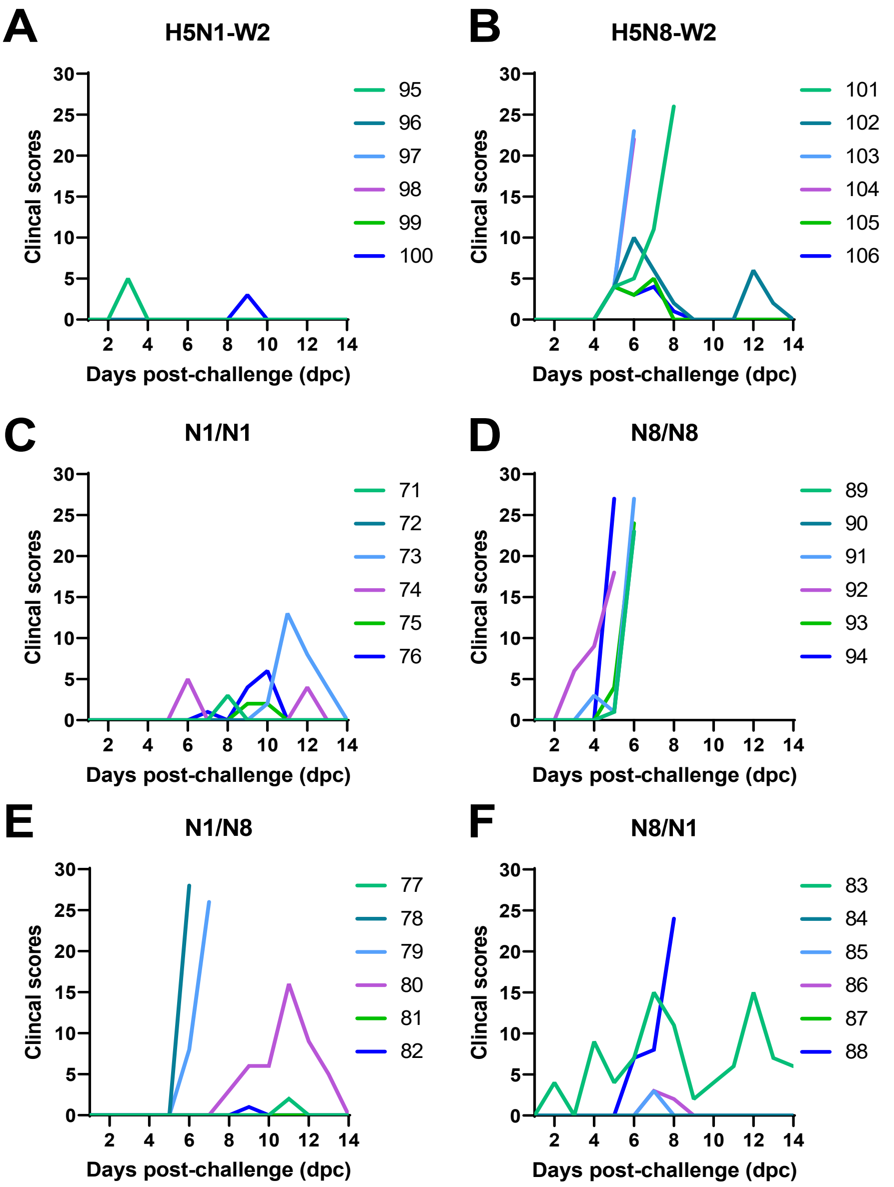

**Fig. S2. Clinical scores following infection with H5N1-W2 or H5N8-W2 with or without prior exposure to LPAIV.**

Individual clinical scores exhibited per timepoint following infection with HPAIV H5N1-W2 (**A**) or H5N8-W2 (**B**) without prior exposure, or in birds which had previously been exposed to a homologous (N1/N1 (**C**) and N8/N8 (**D**)), or heterologous (N1/N8 (**E**) and N8/N1 (**F**)) subtype of de-engineered H5N1-W1 or H5N8-W1 AIVs. Ducks were scored for severity of clinical scores a minimum of twice daily during the study. Each duck's total daily score has been plotted for 1-14dpc.

**Fig. S3**

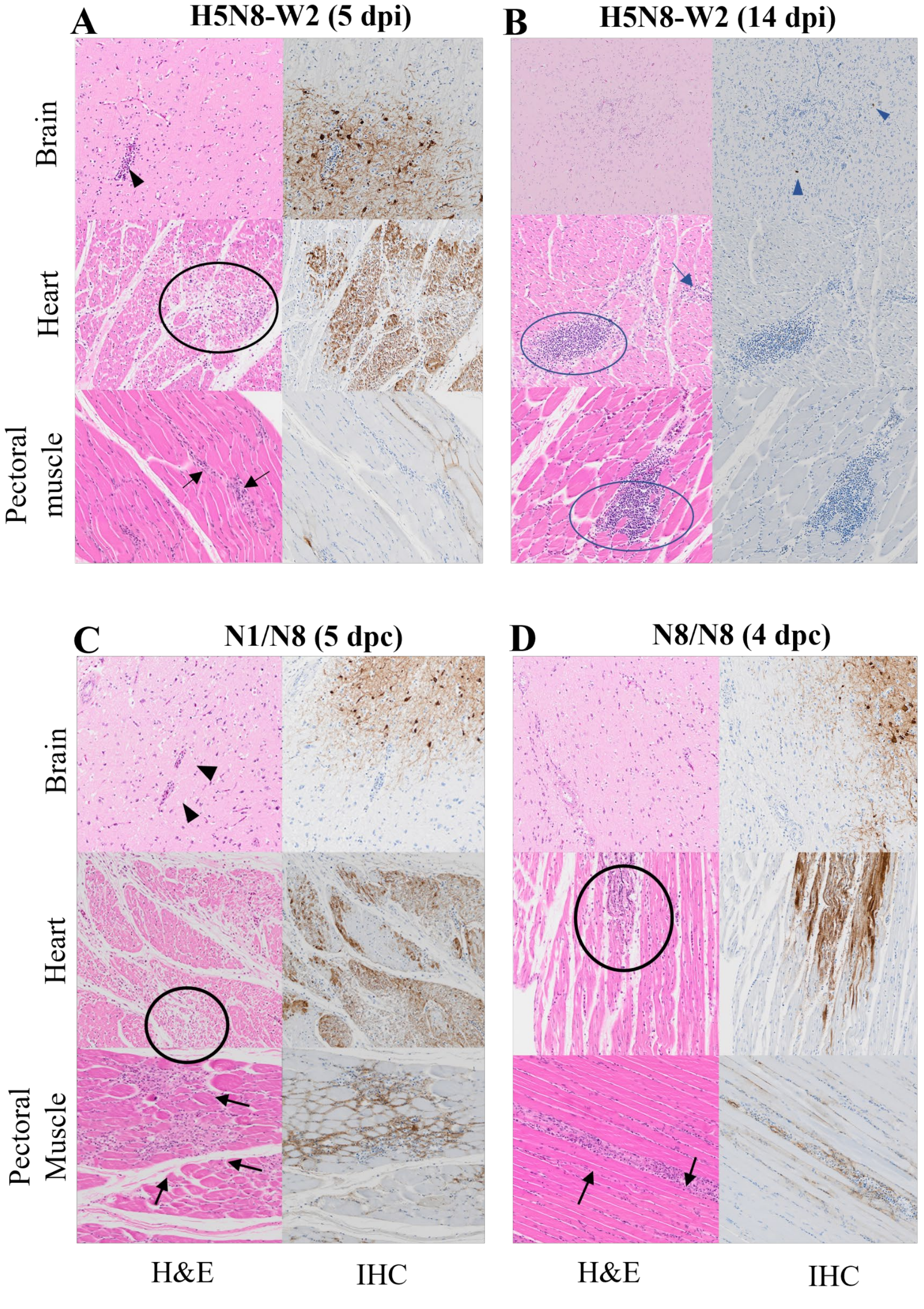

**Fig. S3.** Representative histopathology (H&E) and immunohistochemistry (using an anti-influenza A nucleoprotein antibody staining brown) demonstrating comparison of changes in heart, muscle, and brain from naïve ducks infected with H5N8-W2 at 5 dpi (**A**) and 14 dpi (**B**), and those inoculated with SB-H5N1-W1 followed by challenge with H5N8-W2 (N1/N8) at 5dpc (**C** ), with SB-H5N8-W1, followed by challenge with H5N8-W2 (N8/N8) at 4 dpc (**D**). Acute lesions seen at 4-5 dpi (**A**) or dpc (**C** and **D**) were multifocal acute nonsuppurative encephalitis with mild lymphohistiocytic perivascular cuffing (arrowheads) and abundant immunolabelling in neurons in the brain, myocardial degeneration/necrosis (circles), and myositis, with viral antigen seen in cardiomyocytes in the heart and myocytes in pectoral muscle. Chronic changes at 14dpi (**B**) show minimal multifocal gliosis in the brain and multifocal lymphoid hyperplasia and mild lymphocytic inflammation in heart and muscle, with minimal viral antigen in neurons and macrophages representing viral clearance. Limited lesions were observed in N8/N1 ducks; no samples from N1/N1 were available. Magnification: x20. Days post infection (dpi), days post challenge (dpc).

### Fig. S4

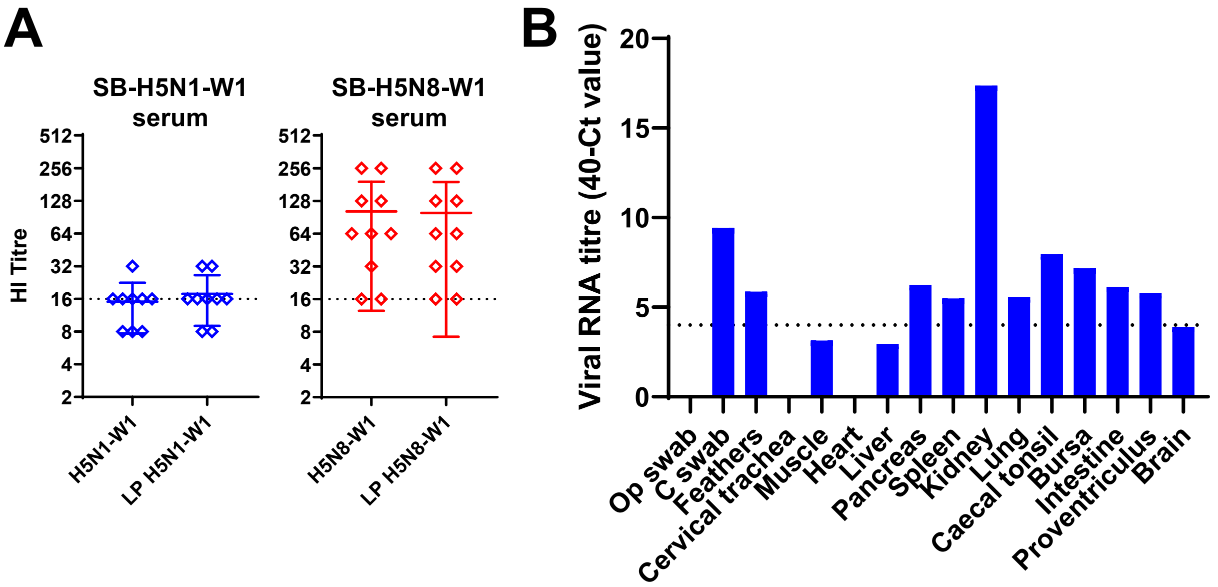

**C**

| | -9 | -8 | -7 | -6 | -5 | -4 | -3 | -2 | -1 | 0 | 1 | 2 |
|---|---|---|---|---|---|---|---|---|---|---|---|---|
| HPAIV H5N1-W1 | P | L | R | E | K | R | R | K | R | G | L | F |
| HPAIV H5N8-W1 | P | L | R | E | K | R | R | K | R | G | L | F |
| SB-H5N1-W1 | P | L | G | E | - | - | - | T | R | G | L | F |
| SB-H5N8-W1 | P | L | G | E | - | - | - | T | R | G | L | F |
| SB-H5N1-W1 (IVPI kidney) | P | L | G | E | - | - | - | T | R | G | L | F |

**Fig. S4 Outcomes from intravenous pathogenicity index (IVPI) of H5N1-W1 and H5N8-W1 LPAIV.**

**A.** Blood collected from nine (SB-H5N1-W1) and 10 (SB-H5N1-W1) chickens inoculated for an IVPI and then tested by HI against homologous wt HPAIV and de-engineered AIV antigens. **B.** Viral RNA detection in organs of one chicken which was found dead at 6 days post-inoculation with H5N1-W1, during the IVPI determination. **C.** Sequence comparison of the cleavage sites of the wild type (wt) HPAIV and de-engineered SB-H5N1-W1 and SB-H5N8-W1, as well as virus found in the kidney sample of the one chicken from **B**.
